# TMRCA is observable from the branch lengths of a coalescent

**DOI:** 10.64898/2026.07.22.740130

**Authors:** Fabian Freund, Daniel Teofilov, Émilien Joly, Arno Siri-Jégousse

**Affiliations:** University of Leicester, United Kingdom; CIMAT, Guanajuato, Mexico; IIMAS, UNAM, Ciudad de México, México

**Keywords:** ancestral recombination graphs, TMRCA, sequential Markovian coalescent, multiple merger coalescents, branch lengths

## Abstract

The height of ancestral trees, in other words the time to the most recent common ancestor in a genealogical tree, is an informative statistic in population genomics and used in contexts of genealogical dating, demographic history inference, and assessing signals of selection. Exploiting a general pathwise identity between the height of a tree and its branch lengths, we revisit Zeng et al.’s mutation rate estimator to develop a model-free estimator for the product of the expected height of genealogical trees with the scaled mutation rate, both for single loci and genomic regions. This estimator is a linear function of the site frequency spectrum of a sample, i.e., of the distribution of allele counts at all segregating sites across the locus or genomic region and thus both observable and computationally cheap. We show that, under the infinite sites model of mutation, our estimator is unbiased and, for two genome-wide models of ancestries (sequential Markovian coalescents and common-pedigree ancestries), consistent. Furthermore, we show via simulation that our estimator performs smaller errors than averaging over reconstructed ancestral tree heights (extracted from reconstructed ancestral recombination graphs) when considering genome properties similar to human genomes across large genomic regions. We then revisit a publicly available genomic dataset of dogs and wolves and compare our method with extracting TMRCAs from ancestral recombination graph reconstructions.

## 1 Introduction

Tree-valued stochastic processes which describe the genealogies of regions across the genome, the ancestry of full chromosomes, have become staple tools in inference methods in population genomics [36, 43] - usually referred to as ancestral recombination graphs (ARGs). Their value comes from their computational efficiency and their ability to link observable genomic diversity with the underlying evolutionary history, which is reflected in the genealogical trees. Multiple inference techniques do not use the full tree sequence but use features of that sequence for inference. A classical example is the PSMC method of [37], which employs a Hidden Markov Model based on the profile of genealogical tree heights (genetically, the time to the most recent ancestor of the two sequence parts, which we abbreviate as TMRCA) across the genome to infer how population sizes of the sampled population have changed. Further demography reconstruction methods also use pairwise or sample-wise TMRCAs as latent variables [58, 57]. Generally, the TMRCA of a genomic region and how it changes across genomic regions due to recombination contains information about various genomic and demographic factors that affected the sampled population, including selection [43, 25, 6]. More fundamentally, the TMRCA of a sample can provide information about the age of the population - at least for some standard population genomic ancestry models [52, 17]. Thus, it is no wonder that understanding the mathematical properties of TMRCAs is an active area of research [60, 64, 32, 65].

The TMRCA at a single genomic region or across the genome cannot be directly observed - we only “see” the genetic signal of mutations that change the sequences in the sample. In the mathematical model for genomic sequences, the mutations arise as random points across the genealogical trees. Under the standard infinite sites model typically used in applications for whole-genome SNP data, a mutation on a branch of the genealogy that ends in *i* leaves changes the genomic type at a (random) nucleotide site for *i* sequences/individuals in the genomic area described by the genealogical tree to a different type (and each mutation hits a different site, mathematically we can assume a continuous genome). Thus, while we have a straightforward connection between the number *ξ*_*i*_ of mutations appearing in *i* sequences in the sample with the sum *L*_*i*_ of the lengths of all branches in the genealogical tree ancestral to exactly *i* leaves, the connection of genomic data with the TMRCA of a sample of size *n >* 2 is not as simple. However, trees are rather regulated mathematical objects, and this has been used in [56] to establish a connection between the TMRCAs of genealogical trees and the branch lengths *L*_1_, …, *L*_*n*−1_, in expectation.

Here, we are assessing the connection between TMRCAs and the spectrum of branch lengths *L*_1_, …, *L*_*n*−1_ in the other direction. We assess how we can express the height of the genealogical tree in terms of the branch lengths *L*_1_, …, *L*_*n*−1_. We will use this to formulate an estimation method of expected genealogical tree heights, scaled by a mutation parameter, based on the site frequency spectrum (*ξ*_1_, …, *ξ*_*n*−1_), the latter being a statistic we can compute from sequencing data. We will then assess the mathematical properties of this estimator both for single genomic regions (without recombination, ancestry described by a single genealogical tree), for small regions (described by a small number of trees) and across the genome (the latter in a mathematically idealized setting of genome length tending to infinity). In order to do this, we will need to establish general mathematical properties of different models for genealogies across a genome.

Finally, we will assess, via simulation of genomic data under realistic parameter assumptions in different evolutionary settings, how big the error of our method is and compare this with estimations based on current reconstruction methods for genealogies. We will then apply our method to a data set from [6] where we reconstruct (expected) TMRCAs from a sequenced chromosome from a combined sample of dogs and wolves in a sliding windows approach, and compare our method to extracted TMRCAs from genealogy (ARG) reconstruction methods.

## 2 A model-free relation for genealogical trees

Denote the time to the most recent common ancestor of a sample of size *n* by *TMRCA*_*n*_ and the time spent in the genealogical tree in a lineage having *i* sampled descendants by *L*_*i*_, *i* ∈ {1,…, *n* − 1}. The sample can be haploid individuals or single genomic strains - we will refer to the sampled units as individuals in either case for simplicity. We assume here that the whole sample was collected at the same time. In the sequel, *L*_*i*_ will be referred to as the branch length of order *i*. See Figure 1 for an illustration. *TMRCA*_*n*_ can be obtained from the branch lengths vector.

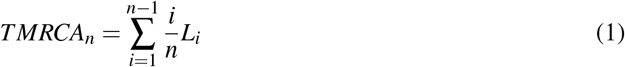

**Figure 1:**
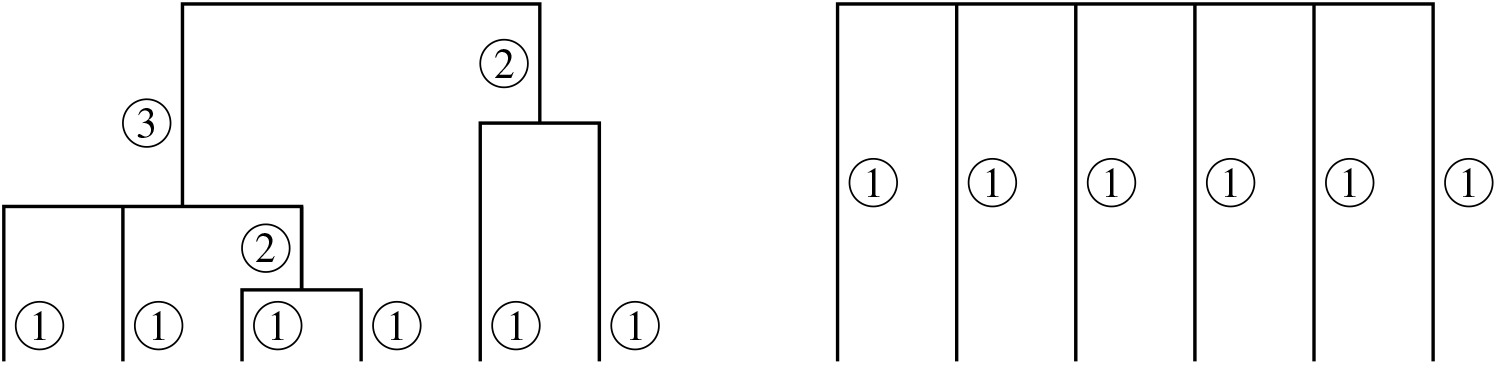
An illustration of relation (1) for *n* = 6. Left: a genealogical tree starting from 6 individuals. Inside the circles appear the order of the lineages, i.e. their number of descendants in the sample. The length *L*_*i*_ is given by the sum of the lengths of the branches labeled by *i*. Right: tree obtained by replacing lineages of order *i* by *i* lineages of order 1. Its total length is given by *nTMRCA*_*n*_.

This can be understood thanks to Figure 1, replacing each branch contributing to *L*_*i*_ with *i* copies of itself leads to a star-shaped tree where all branches only merge at the root. For formal mathematical proofs for results in this and all other sections, see the Appendix.

This relation is a general statement about ultrametric mathematical trees with real-valued branch lengths. Here, ultrametricity corresponds to equal sampling times across each individual. There is an analogous argument if the tree is not ultrametric, i.e. we look at different sampling times of the leaves in case of genealogical trees. If so, we have to understand *TMRCA*_*n*_ as the time between the most recent common ancestor of the sample and the present time (time 0, the most recent sampling time). This is because individuals *k* have been sampled at different times *t*_*k*_ in the past, and then have distinct distances to the MRCA. Thus, formula (1) modifies to

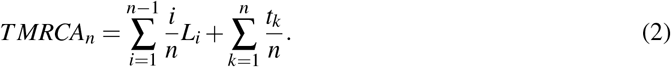

This is easily seen from Figure 2. We dedicate a complete study of this case in Appendix 10.2.

**Figure 2:**
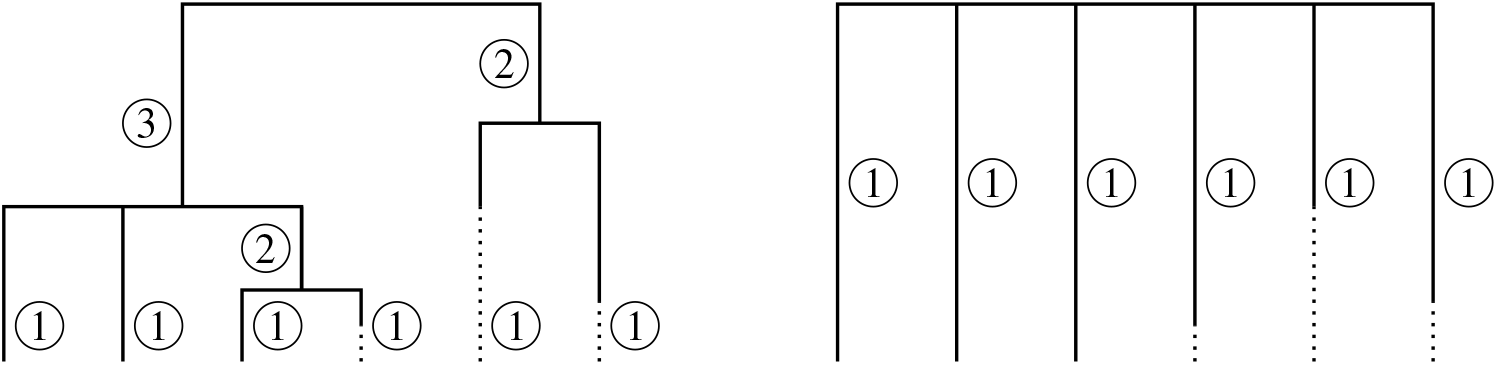
An illustration of the relation when serial samples are performed. Adding the dotted lengths (*t*_*k*_’s in Eq. 2) leads to ultrametric trees for which Eq. (1) holds.

The relation can be generalized to other types of genealogical structures. As an example, consider the ancestral recombination graph (ARG) with two loci and sample size *n*. We define this not as a single graph, but equivalently by considering two ancestral trees at the two different loci, see also Appendix 10. The time to the most recent common ancestor at both loci 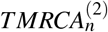 is given by

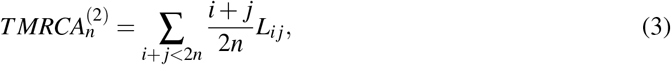

where *L*_*i j*_ is the branch length in the ARG with *i* ≥ 0 descendants at the locus *a* and *j* ≥ 0 descendants at the locus *b*, before the time to the most recent common ancestor. With a straightforward generalization of notation, we get for the ARG with *k* loci and a sample size *n*,

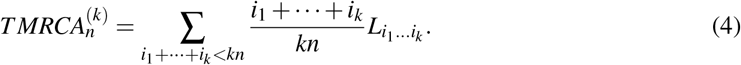

## 3 Coalescents and first applications

Although relation (1) works for any tree model, we will mainly consider random coalescent models in the sequel, namely Ξ-coalescents [54]. Coalescents provide a wide class of possible genealogical trees, which most commonly used tree model is Kingman’s coalescent, the standard genealogical model for neutral evolution [59].

Mathematically, Ξ-coalescents are continuous-time Markov chains that track the lineages of a sample of size *n* backward in time. A coalescent starts with *n* separated lineages and allows them to merge over time, increasing their size, until reaching one lineage of size *n* (the MRCA). A Ξ-coalescent can have simultaneous and multiple mergers, see Figure 3. Its Markov transition matrix, describing at which rate and how lineages merge can be obtained explicitly as a function of a finite measure Ξ on the simplex Δ = {(*x*_*i*_)_*i*_|*x*_*i*_ ≥ *x*_2_ ≥ … ≥ 0, ∑_*i*_ *x*_*i*_ ≤ 1}. Examples of Ξ-coalescents are the symmetric coalescent [22, 21], representing limit genealogies of populations under recurrent bottlenecks, or Beta-Xi-coalescents [7, 9], modeling genealogies of diploid populations under reproduction sweepstakes.

**Figure 3:**
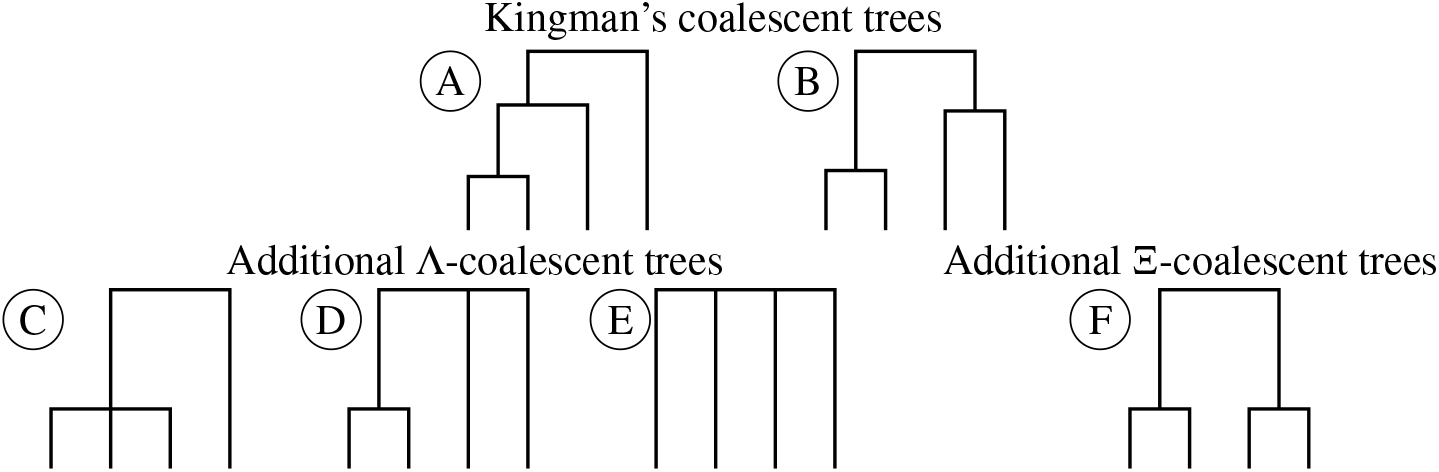
The six possible Ξ-coalescent topologies for four sequences.

An important subclass of coalescents called Λ-coalescents, are those where no simultaneous merger is allowed, see Figure 3. Among those multiple merger coalescents, the celebrated family of Beta-coalescents provides genealogical models for populations with skewed offspring distributions, but also has been discussed as a model for range expansions [8]. At both sides of the range, the Kingman coalescent models neutral evolution and the Bolthausen-Sznitman coalescent models rapidly adapting evolution (e.g., [13, 41]) and other selection modes [53]. For a variety of species, different Ξ- and Λ-coalescents have shown to fit well to genomic diversity data, and especially better than purely bifurcating genealogy models [28, 45, 4, 18, 40, 26, 3, 20].

### Applications to classical coalescents

We are able to recover the expected TMRCA when the expected partial lengths are known. This is the case for Kingman and Bolthausen-Sznitman cases.

For the Kingman coalescent, E[*L*_*i*_] = 2*/i* and does not depend on *n*, so we recover the classical result

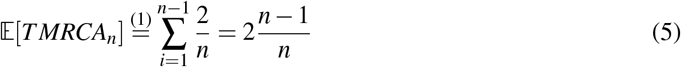

For the Bolthausen-Sznitman coalescent with initial sample size *n*, we know from [31] that

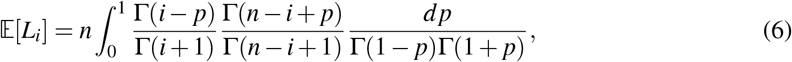

and we can obtain a new formula for the expected TMRCA.

#### Proposition 3.1.

*Set ĉ*_0_ = 1 *and, for k* ≥ 1, 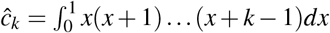 *be the Cauchy numbers of the second kind. Then*

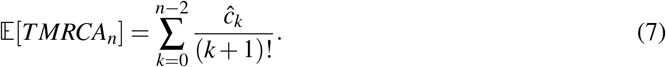

*holds for the Bolthausen-Sznitman coalescent*.

## 4 Genetical interpretation - single locus

### Mutations model and the observable statistics

The pathwise identity (1) of the weighted branch lengths sum and the TMRCA can also be described in terms of genomic data. First, we consider a single locus without recombination sampled in *n* individuals or genomic strains. If the coalescent tree is representing the genealogy of a sample of size *n*, the mutations seen in the sample can be represented as produced by a homogeneous Poisson process with rate *θ* on its branches. Under the infinite-sites model, each mutation hits a different site in the genomic region described, and each individual inherits all mutations on the path in the coalescent from the root (MRCA) to the leaf representing this individual.

In this setup, a natural analogon of the branch lengths *L*_*i*_ is the number of mutations *ξ*_*i*_ on these branches - Poisson distributed given *L*_*i*_. The vector (*ξ*_1_, …, *ξ*_*n*−1_) is called the site frequency spectrum. It is one of the most prominent measures of genetic diversity in population genetics and is a classical statistic for parameter estimations and model selection [19]. In particular, we have E[*ξ*_*i*_] = *θ* E[*L*_*i*_]. Thus, (1) leads to a first moment estimator of *θ TMRCA*_*n*_. With

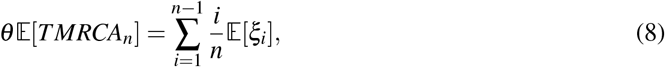

we can provide the estimator

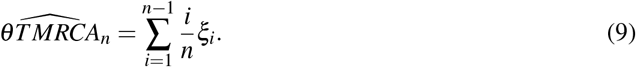

### The Zeng estimator revisited

Interestingly, the same equation, when plugging in the expected value for an underlying Kingman’s coalescent genealogy, leads to Zeng’s mutation rate estimator, and a basis for deviation from neutral evolution test, introduced in [63]. In more detail, we have

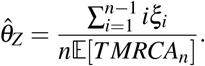

Under the Kingman coalescent model with E[*TMRCA*_*n*_] given by Eq. 5, we recover the unbiased estimator for *θ* from [63]. Recall that the mutation rate *θ* is replaced by *θ/*2 in the referenced paper, canceling the factor 2 below:

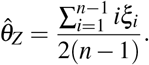

We can also compute generalized versions under other coalescent models. For the Bolthausen-Sznitman coalescent, Proposition 3.1 provides an unbiased estimator for *θ* as

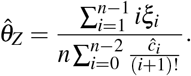

In general, under any Markovian coalescent model, including inhomogeneous ones such as exponential growth or bottlenecks, the expectation can be obtained via computational techniques based on phase-type theory, see [60, 24]. As commented in [63], this estimator gives more weight on ancient mutations compared to other *θ* estimators as nucleotide diversity or Watterson’s estimator.

### Estimating the TMRCA

For practical purposes, we are usually less interested in estimating *θ* E[*TMRCA*_*n*_] than estimating E[*TMRCA*_*n*_], as the latter would allow us to date the sampled population, while the former just allows comparisons of the TMRCAs between different genomic regions/chromosomes or samples from different populations within the same species (technically, we can compare wherever *θ* is identical). However, if we have a precise estimator for the mutation rate *µ*^∗^ for the genomic region described by the genealogical tree on any timescale (by generation, by year,…), e.g. from a mutation accumulation experiment or from generation-by-generation comparisons within a known pedigree, 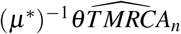 then provides an unbiased estimator of E[*TMRCA*_*n*_] in the time unit of the mutation rate estimator. Usually, we will have a per basepair, per time unit mutation rate estimate *µ*. In this case, if mutation rates can be assumed to be stable across the genomic region of length *l* basepairs, *µ*^∗^ = *lµ*.

### The renormalized statistics

In genetic terms, 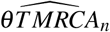 is summing up the allele frequencies of the mutations appearing in the sample described by the genealogical tree. A related and directly interpretable object would be the average allele frequency, thus dividing by 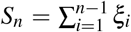. The expected value of 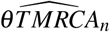 is *θ* E[*TMRCA*_*n*_] and 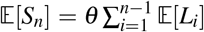. Heuristically, this means that the average allele frequency compares the TMRCA with the total length of the tree. Mathematically precise, we can say the following: the expected value of the average allele frequency equals

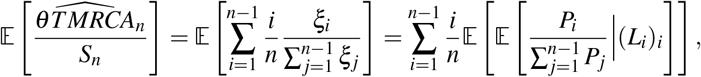

where the *P*_*j*_’s are independent Poisson r.v. with rate *θ L*_*j*_. Using elementary properties of Poisson random variables and their ratios and the convention that, with no mutations, the value of the ratio is 0, we observe 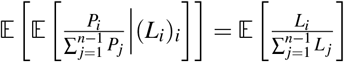. Thus, we can compute

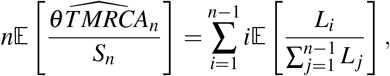

which is an expected value by the probability measure defined via the expected values on the right hand side. In other words, the average allele frequency is the average number of samples a random ancestral point (a point randomly placed on the genealogy) has as descendants. This does not depend on the mutation rate *θ*.

## 5 Genetical interpretation - genome-wide

Our interest lies in using (1), more precisely its first-order observable approximation (9), as a moment estimator for *θ TMRCA*_*n*_. Our description holds true for mutations for the case of a single genealogy. Apart from extremely clonal species as *Mycobacterium tuberculosis*, genomes will trace back different genealogies in different genomic regions due to recombination. So what does our estimator from (9) measure in the case of multiple genealogies, e.g. in a diploid, recombining species? If *θ*_*b*_ is the mutation rate per base pair (on the timescale of the genealogical tree), and we assume this to be constant across the genome, each region of length *l* with shared genealogy will then have an effective mutation rate of *θ* = *θ*_*b*_*l* on the genealogy. However, this genealogy may change at each recombination point, as recombination leads to a change in ancestry of the genomic material from different sides of the recombination event. Thus, the ancestral process across the genome consists of a series of trees, changing through recombination. These local genealogies will still be correlated, at least over short genomic distances, as recombination changes only certain branches in the genealogical tree. Thus, a genomic region of length *l* may not estimate *θ TMRCA*_*n*_, but a weighted sum of different 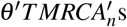. That being said, not every recombination event will change the MRCA, especially for large samples for which the probability that a recombination point hits near leaves of the genealogical tree gets bigger [52, 17].

For these reasons, we may not be able to measure the TMRCA of a given segment by our estimator. However, if mutation rates are constant across basepairs, 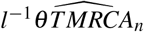 will estimate an average TMRCA, weighted by the “sojourn genomic region” whose genealogy is given by a specific tree. We will now analyze the mathematical and numerical properties of the estimator computed on large genomic regions/genome-wide for the Sequential Markovian Coalescent (SMC) [39] and its extension to Beta-coalescents (SM*β* C) from [33] and other multiple merger genealogies, as well as for the multiple-merger genealogy model with recombination from `msprime` [5, 34]. We can show that we expect the genome-wide TMRCA to be close to the expected TMRCA of the underlying one-locus coalescent tree.

### 5.1 Consistency for genome size *l* → ∞ due to ergodicity of sequential Markovian coalescents

As we consider an estimator, a natural question is whether it is consistent, i.e. that it approximates the value it estimates with an error that vanishes with ever increasing data. For genealogies across the genome, this means we want to allow genome size *l* → ∞. We will also assume that the mutation rate is constant across the genome. Let *θ*_*b*_ be the mutation rate per basepair

First, we recall the definition of the SMC and the SM*β* C, which allow precise mathematical treatment. Moreover, we extend the definition of the SM*β* C proposed in [33] to the more general case of allowing local Ξ-coalescent genealogies, the Sequential Markovian Ξ-Coalescent (SMΞC) for any underlying Ξ-*n*-coalescent. This class includes the standard SMC (with underlying Kingman coalescent). For all of these models, we can establish that 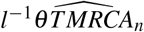 is a consistent estimator for *θ*_*b*_E[*TMRCA*_*n*_], where E[*TMRCA*_*n*_] is taken from the model without recombination. This follows from the ergodic properties of the ancestral tree processes across the genome.

The Sequential Markovian Ξ-coalescent extends the Ξ-coalescent by adding a recombination mechanism which changes the trees across the genome. Recall from Section 3 that the coalescence rates of a Ξ-coalescent can be expressed in terms of the finite measure Ξ on Δ = {(*x*_*i*_)_*i*_|*x*_*i*_ ≥ *x*_2_ ≥ … ≥ 0, ∑_*i*_ *x*_*i*_ ≤ 1}.

To construct these sequential Markovian coalescents, start with a genealogical tree drawn from an underlying distribution. The recombination event happens at a uniform position across all branches of the current genealogy, removes a lineage from the genealogy from this time point further towards the root (MRCA) of the genealogy, and “reglues” this lineage to the remaining tree in a way that preserves the underlying coalescent properties.

For any Ξ-coalescent with Ξ((0, 0, …)) *>* 0 and Ξ(Δ \ (0, 0, …)) *>* 0, i.e. which allows for multiple mergers but does also have purely binary mergers that cannot increase to multiple mergers when sample size increases, the construction will have an additional step, which we will add in quadratic brackets [MMC+KM:…].

We will now describe how to construct the Sequential Markovian Ξ-coalescent of a sample of *n* initial lineages/individuals and recombination with rate *ρ* ≥ 0.

1. Consider a sample of *n* individuals 1, …, *n*, all of them having an infinite genome [0, ∞).
2. Start with a tree 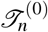 with total branch length 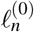 at genomic position 0, generated as a realization of the underlying Ξ-coalescent model for a single locus. [MMC+KM: mark all purely binary mergers]
3. A recombination events appears at a random genomic location, by a changed ancestral relation of the next genomic region compared to the current.
  - The next recombination event happens at genomic position *G*_0_, an exponential r.v. of parameter 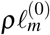.
  - At this genomic position, the ancestral tree changes at a recombination point picked uniformly at random across the current ancestral tree 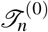.
  To achieve this, we use a Poisson process construction. Let (*x*_*i*_, *t*_*i*_, *k*_*i*_) be the ranked atoms (according to their first coordinate) of a Poisson point process on [0, ∞) ×[0, ∞) ×[*n*] with intensity *ρdx* ⊗ *dt* ⊗ *dk*, where the first coordinate describes genomic position, the second describes time in the past and the third describes the label of the lineage where the recombination point would occur (we label each ancestral lineage by the smallest index of the sampled individual). If at time *t*_1_, *k*_1_ is the label of any present ancestral lineage of 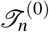, a recombination event occurs on this branch of 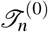 at this time, at genomic position *x*_1_. If not, consider the next atom (*x*_2_, *t*_2_, *k*_2_) and repeat the procedure, see Figure 4.
4. Produce a new tree 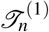 from 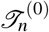 by freeing and regluing the lineage at the recombination point. See Figure 5 for an illustration. This tree is the ancestry for the region starting at *G*_0_.
  - Cut the branch of 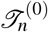 at the uniformly chosen point and remove the branch extending towards the root from the tree. This lineage is “freed”.
  - Recoalesce following the approach from [14], which will be extended below to Ξ-coalescents. This lineage is “reglued”.
5. Repeat the procedure on the mew tree 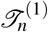 with total branch length 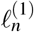, using the following atoms of the Poisson point process to create a new tree 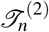 at the genomic position *G*_0_ + *G*_1_ where *G*_1_ is an exponential r.v. of parameter 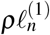.
6. Repeat this *ad infinitum*

**Figure 4:**
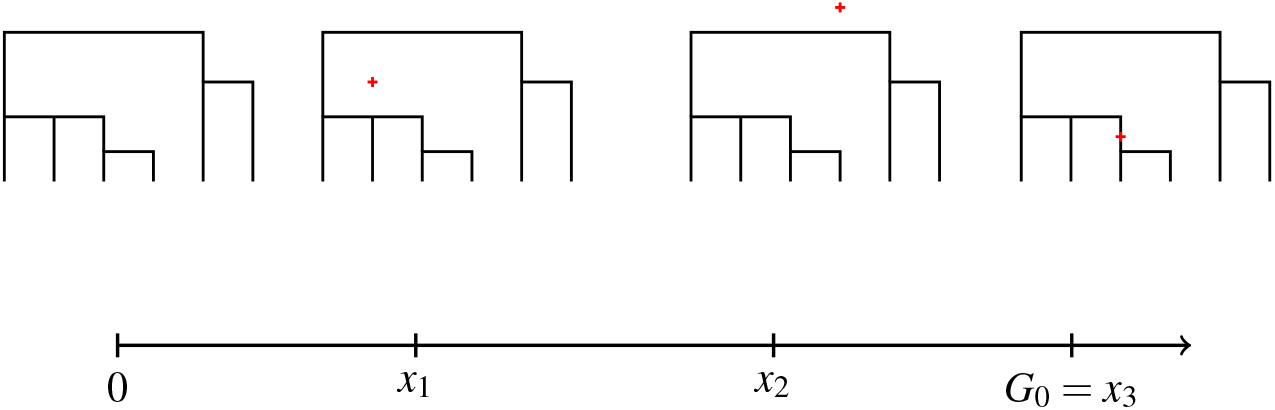
An illustration of the recombination point picking in the SMΞC. At genomic positions *x*_1_ and *x*_2_ the times and labels (*t*_1_, *k*_1_) and (*t*_2_, *k*_2_) do not intersect with the tree. At genomic position *x*_3_ the time and label (*t*_3_, *k*_3_) intersects with the tree so that a freeing and regluing procedure can occur at this position..

**Figure 5:**
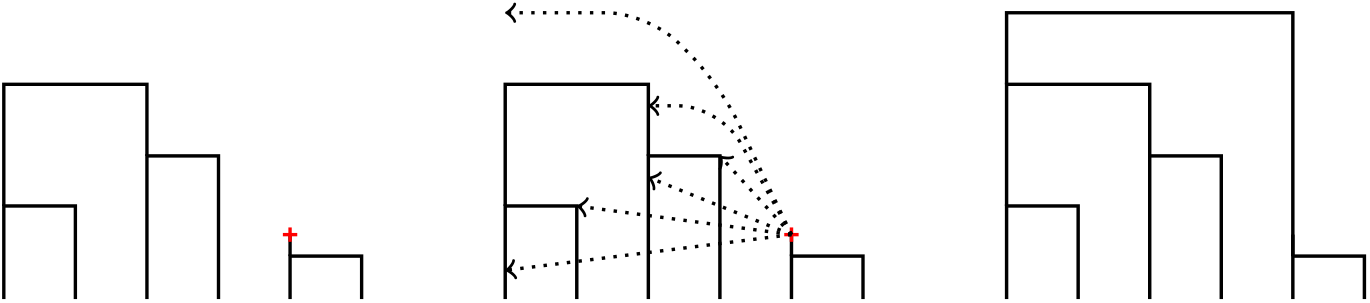
An illustration of the freeing and regluing procedure. Left: the ancestral lineage cut at the recombination point is freed. Center: some options for the freed lineage to reglue. Right: the new tree, after regluing the freed lineage

Now we will describe in detail the regluing operation: Recall from [54] that the Ξ-coalescent, in a current state with *b* ancestral lineages, has infinitesimal rates *λ* (*b*; *k*_1_, …*k*_*l*_; *s*) with *k*_*i*_ ≥ 2 and 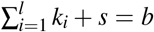. These rates describe transitioning the state with *b* lineages by merging simultaneously specific sets of *k*_1_, …, *k*_*l*_ of the present *b* lineages into *l* separate lineages, while keeping *s* lineages unmerged. Following the argument in [14, Sect. 5], we end up with a Ξ-coalescent by construction if we let the freed lineage re-coalesce with the same distributional properties as conditioning one lineage of a *b*-coalescent on the behaviour of *b* − 1 lineages, which directly follows from the rate consistency relationship ([54, Eq. 23]). So a freed lineage (at time *t*) reglues, conditional on the remaining tree, from *t* onwards, with one or more of the non-freed *b* − 1 lineages, in the following way.

- Merge with any present lineage at a time where no other coalescence event is present in the remaining tree at rate (*b* − 1)*λ* (*b*; 2; *b* − 2). [MMC+KM: this rate splits into rate (*b* − 1)*c* with *c* = Ξ((0, 0, …)) and into the binary merger rate from Ξ^′^. Mark purely binary mergers.].
- Join the specific existing merger of *k*_*i*_ lineages within an existing (*b* − 1, *k*_1_, …, *k*_*l*_; *s*)-merger of the remaining lineages with probability 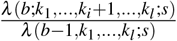 or, if *s >* 0, add an additional binary merger to this coalescence event with probability 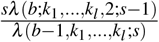 [purely binary mergers cannot be joined at all]

#### Remark 5.1

(Main differences with the original SMC). *Although the framework may look different, most of the properties of the SMC [39] and its modification to Beta-coalescents [33] transfer to the* Ξ *case. In contrast to the original definitions, who use a finite genome* [0, 1] *and a recombination rate ρ/*2, *the genome is now infinite and the recombination rate is now ρ*.

We will now establish an ergodic result for the SMΞC. Note that this also works for its Kingman and Λ-coalescent version. Let 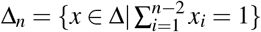 for *n* ≥ 3.

#### Theorem 5.1.

*Consider any* Ξ*-coalescent for n* = 2 *or satisfying* Ξ(Δ_*n*_) *<* Ξ(Δ) *for n* ≥ 3. *For the associated SM*Ξ*C, the distribution of the underlying* Ξ*-n-coalescent is a unique invariant distribution. Let TMRCA*_*n*_(*l*^′^) *denote the time to the most recent common ancestor in the SM*Ξ*C at genomic position l*^′^ ∈ [0, *l*), *where l is the genome length. Let TMRCA*_*n*_ *be the time to the most recent common ancestor in the associated (single locus)* Ξ*-n-coalescent. Then*,

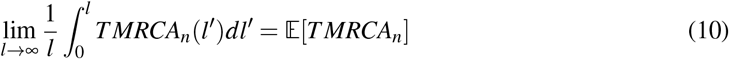

*almost surely and in L*^*p*^, *p* ≥ 1. *The analogous convergence to the single-locus mean holds for any p-integrable function under the invariant distribution, including all entries of the length vector* (*L*_1_, …, *L*_*n*−1_).

The family of Ξ-coalescents that do not have associated ergodic SMΞCs is a small model class. They have no purely binary mergers. Their first merger (while *n* lineages are present) has to be a multiple merger or consist of at least two simultaneous mergers, and this cannot be fully dissolved by freeing and regluing - thus all trees in the genome have a coalescence event at this timepoint. The most prominent coalescent in this class is the star-shaped coalescent, which merges all lineages after an exponential time, and any freed lineage just reglues to the same merger again. Observe that the class of symmetric coalescents [22], that models recurrent bottlenecks phenomena in the evolution, satisfies the assumption Ξ(Δ_*n*_) *<* Ξ(Δ) as long as the size of the remaining population after the bottleneck can be larger than *n* − 2.

The ergodic theorem for the SMΞC allows us to derive genome-wide consistency (for genome length tending to infinity) of our estimator from (9), whose genome-wide version is divided by the genome length *l*, i.e.

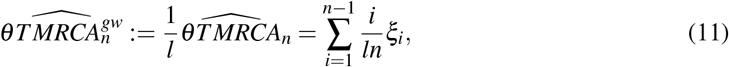

with *ξ*_*i*_ the *i*th entry of the genome-wide SFS.

#### Corollary 5.1.

*For any SM*Ξ*C*, 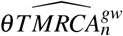 *is a consistent estimator of θ* E[*TMRCA*], *i*.*e*.

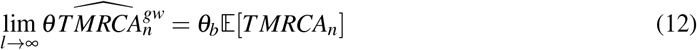

*in probability*.

As in the single-locus case, if we have a precise per basepair estimate *µ* of the mutation rate on any timescale (generations, years, days, …), 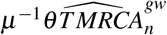 will be a consistent and unbiased estimator of E[*TMRCA*_*n*_] on the same timescale.

### 5.2 Sequential Markovian coalescent in the presence of genome-wide effects

While the SMC model is well-established as an approximate genome-wide model when Kingman’s coalescent is the local (one-locus) model [47, 38], its multiple merger version so far has only been used as a proxy and easier-to-handle model in [33] for parameter inference of a different genome-wide multiple merger coalescent model, following [34], which models that multiple merger events should affect all genomic regions. The latter model is the basis of the implementation of multiple merger coalescents with recombination in the coalescent simulator `msprime` [5]. We are tempted to use SMΞC as a conceptual model for genealogies for genome-wide data in reproduction models whose one-locus genealogies are in the domain of attraction of Ξ-coalescents. However, a general issue with this use is that, if the multiple mergers are due to large families appearing over small timespans, this should affect all regions in the genome, which is not a property of the SMΞC. This has been recently discussed in context of the shared pedigree for these regions (e.g., [2], [15],[42]). Many mechanisms we discussed earlier which are linked to the appearance of multiple mergers at single loci should lead to an (potential) effect in all genomic regions, e.g. extreme bottlenecks, sweepstake reproductions and range expansions. However, we do conjecture that selection-driven multiple merger events which can be “escaped” via recombination, as in [16] could be reasonably well approximated by a suitable member of the SMΞC process class.

Having discussed this leads to a natural new question. Can we also formulate an equivalent result as Theorem 5.1 that covers genome-wide genealogies with shared multiple merger events across all loci, i.e. with a shared pedigree? Indeed, we will show that, conditionally on the mechanism producing these large events (potentially) leading to multiple mergers, an analogous result holds. But first, we need to specify the process generating the genealogical trees across the genome, in the presence of these shared large effect events. We will follow [34] to define this process. Instead of conditioning on the pedigree itself we will condition on the process generating the large family events. This is based on the Poisson construction of the underlying Λ- or Ξ-coalescent models for one locus ([48], [54]), see Appendix 10.4. In the sequel, *c* will denote the rate of purely binary mergers, and *p* = (*t, x*) will be the atoms of the Poisson point process Ψ on [0, ∞) × Δ encoding the multiple mergers. As proposed by [42], we will call this process the Ψ-directed ancestral recombination graph (Ξ-ARG).

The Ψ-directed Ξ-ARG with recombination rate *ρ* is defined via Poisson construction of the corresponding one-locus process, with Ξ = *cδ*_0_ + Ξ^′^ where Ξ^′^((0, 0, …)) = 0. We start at genomic position 0 with a one-locus Ξ-coalescent obtained from the atoms of Ψ. Then, we free lineages as in the SMC/SMΞC, with recombination rate *ρ*. Each freed lineage can merge with a present lineage at the genomic position with rate *c*, and at each Poisson point *p* appearing after the lineage is freed, the freed lineage *k*^′^ generates another instance of its paintbox variable *X*_*k*_′ (*p*) and then merges with other lineages if these generate the same index *i >* 0 with their *X* (*p*)-copy, see Appendix 10.4. All lineages at all genomic positions use paintbox variables based on the common Poisson process Ψ.

Clearly, this definition is the standard SMC if Ξ^′^ = 0 and *c* = 1. If *c* = 0, it is the full ancestral recombination graph where we keep track of all past lineages and can remerge with any lineage ever visited (hence we called it an ARG and not a SMC).

The Ψ-directed Ξ-ARG, conditioned on Ψ = *ψ*, is not directly Markovian, as we can merge at Poisson points where the current tree had no coalescences, but previous trees had a merger, if the variables *X*_*k*_(*p*) of the present lineages lead to this coalescene event. However, we can easily make the (conditioned) process Markovian by not only recording the tree itself but also the underlying (*X*_*k*_(*p*))_*p*∈*ψ*_ of each present lineage *k* - after a merger, the merged lineage uses the variables of the lineage with the smallest index (the lineages at genomic position 0 are indexed by 1 to *n*, each freed lineage gets the next integer not used as an index before). We call this process the Ψ-directed SMΞC, and state a conditional equivalent of Theorem 5.1.

#### Theorem 5.2.

*The* Ψ*-directed SM*Ξ*C, conditioned on a realization of the associated Poisson point process* Ψ = *ψ, is ergodic and* (10) *holds for almost all realizations of the Poisson point process, as well as* (12), *for the conditional mean* E(*TMRCA*_*n*_|Ψ = *ψ*).

### 5.3 Estimation errors for finite genomes

The previous section establishes that, for sufficiently large genomes, our mutation-rate scaled TMRCA estimator provides an unbiased and consistent estimate of the expected *θ* TMRCA. But how large is sufficiently large? Moreover, how does 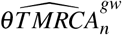 compare to estimating the scaled expected genome-wide TMRCA from reconstructions of the ancestries across the genome (ARG) with state-of-the-art methods? We tested this via simulating a medium-sized to large genomic segment with realistic properties and assessing the performance of 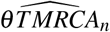 and comparing it against ARG reconstruction with `tsinfer`/`tsdate`. [29, 61].

#### 5.3.1 Simulation approach and results

Using the `msprime` Python library [5], we performed 4,500 simulations of 100 diploid genomes of length 50Mb (50 million basepairs) with per basepair, per generation mutation and recombination rates *µ* = 1.15235 × 10^−8^, *ρ* = 1.29 × 10^−8^ as well as ancestral population size *N*_*anc*_ = 10,000 as indicated for human genomes [1, 35]. A size of 50 Mb is roughly the size of human Chromosome 22, the smallest non-sex chromosome in humans. These 4,500 simulations were evenly divided between simulating three different parameter values alphas of the Beta-Xi coalescent [9, 7] as a genealogy model class for diploid populations. We chose *α* ∈ {1.1, 1.5,, 2}, where *α* = 2 represents the Kingman coalescent, each with per generation exponential growth rates *g* ∈ {0, 0.001, 0.01}.

From the simulated genealogies with mutations, we extracted the site frequency spectrum using the Python library `tskit` [50] to then compute the segment-wide 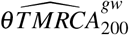. We then computed the mean normalized absolute error (NAE), averaging 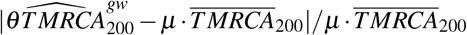, where 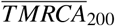 is the average of simulated tree heights across the segments weighted by fraction of the segment where this tree is the genealogy. The tree heights were extracted using `tskit`.

We then compared these errors with the NAEs of the weighted average 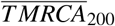 with averaging the heights of reconstructed local genealogies across the segment, weighted by the fraction of total segment length covered by each tree (as genealogy). To reconstruct the genealogies with estimated branch lengths in generations, we used the Python libraries `tsinfer` [30] and `tsdate` [62]. To do this, we exported the simulated sequences from `msprime` as vcf and transformed them to the required input format using the `bio2zarr` library [10]. To estimate the branch length via `tsdate`, we used the true per site, per generation mutation rate used for the simulation. As the data are simulated, we know which allelic state is derived at each SNP, which allows us to perfectly polarise the SNPs and thus the site frequency spectrum. Note that, as NAEs are in units of the true tree heights, the NAEs between a scaled estimator and the analogously scaled TMRCA is identical to the NAE of the unscaled objects. Thus, we can compare the NAEs for 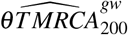 with the NAE of the `tsinfer`/`tsdate`.

Comparing the NAEs, we see that both estimations have small relative errors (mostly below 0.1, i.e. 10% of the true value, for 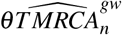), with 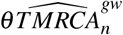 having error distributions closer to 0, see Figure 6. To check for statistical significance we performed a one-sided paired Wilcoxon test for difference in median across the nine cases, all of which came up as statistically significant (*p <* 2.2 × 10^−16^). Interestingly, the distribution of errors, across all different parameters, is more or less centered for 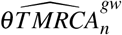 (for 51% of simulations, 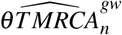 predicts too low TMRCAs), while it is strongly skewed towards overprediction for the extracted heights from the reconstructed genealogies (only for 4.7% of simulations, predictions are too low).

**Figure 6:**
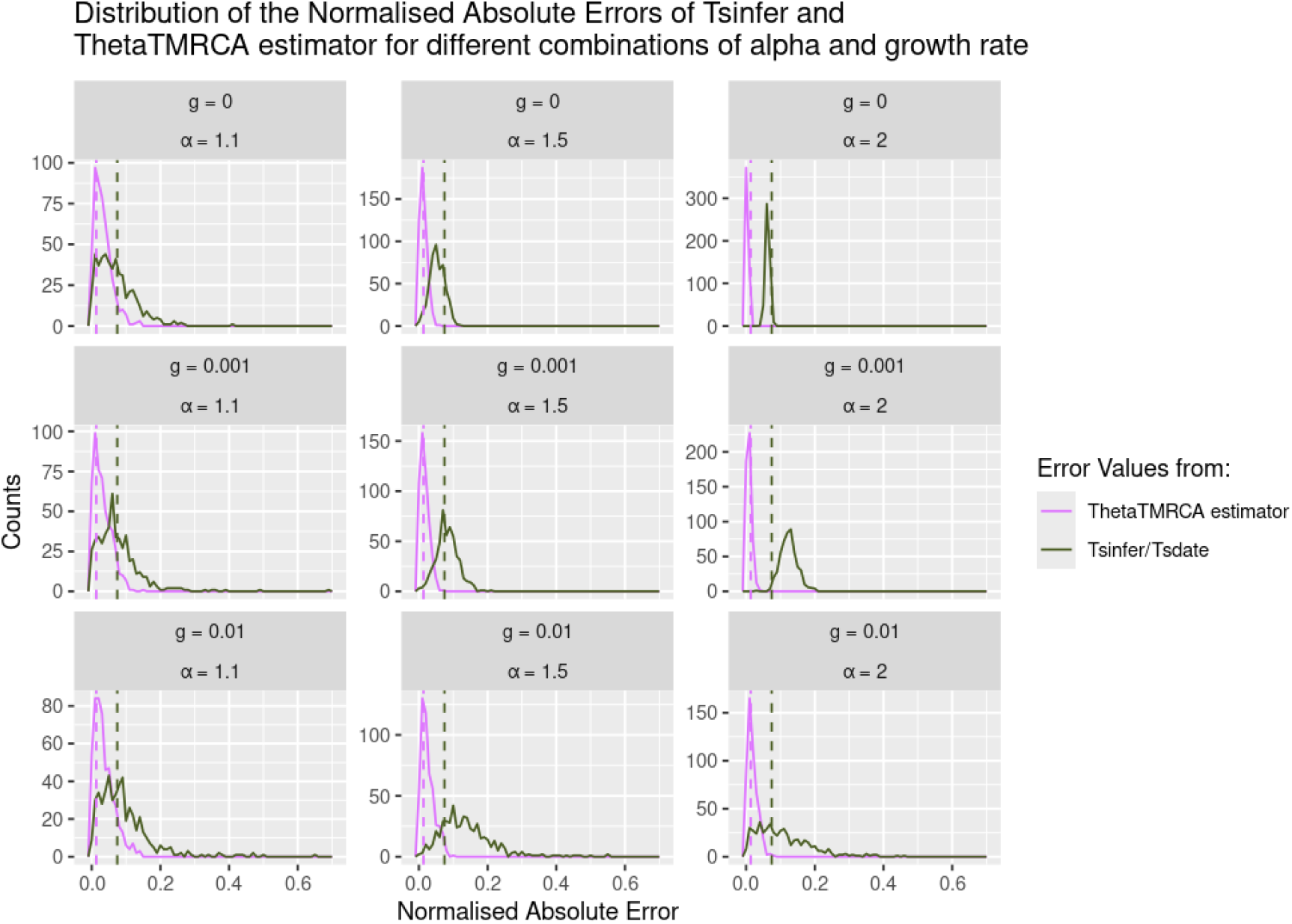
Distributions of normalized absolute errors between estimated and true average tree heights. *α* is the Beta-Ξ-coalescent parameter, *g* is the growth rate per generation. Dashed vertical lines show the distribution medians.

## 6 Application to data: revisiting Chromosome 25 in *Canidae*

In addition to simulated sequences, we also use our *θ* TMRCA estimator on real data to provide TMRCA estimates across regions as a window-based statistic. Strong evidences for selection of the IFT88 gene on dog and wolf chromosome 25 were previously reported by [6]. One of the evidences for this was a visibly smaller value of TMRCA at the gene location measured from reconstructed genealogies across the genome via `relate` [55]. To see how 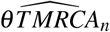 performs on real data, we will again compare the tree heights from reconstructed genealogies with our estimator.

As data, we follow [6] and use genomic (biallelic SNP) sequences of 134 dogs and wolves from Chromosome 25 originally from [49]. We use the view command of `bcftools` first with flag `-r 25` to extract chromosome 25 and then with flags `-v snps -m 2 -M 2` to remove non-biallelic SNPs. Then, in order to extract the samples studied in [6], we use `bcftools` again to subset, excluding the coyotes as the selective sweep is only within dogs and wolves.

We again use `tsinfer/tsdate` package to reconstruct genealogies, for which we need an estimate of the mutation rate for the species considered and to call the derived allele for each SNP. As discussed in [49], we can use a mutation rate *µ*_*dogs*_ = 4 × 10^−9^ for dogs and wolves and an outgroup sequence from the Andean fox to polarize the mutations.

From the reconstructed genealogies, we extracted the TMRCA of every tree and record the TMRCAs as a stepwise constant function across the chromosome. Similarly, we computed the site frequency spectrum from the reconstructed trees via `tskit` using 50,000 uniformly sized windows (length ~1033 bps), then computed 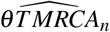 within each window and expressed the TMRCA at each bp via the windowed estimate. We then divide 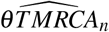 by the mutation rate *µ*_*dogs*_ assumed for dogs and wolves times the length of a window. We compare this with the original reconstruction results using `Relate` [55] from [6], where we extracted TMRCAs of the same sample of dog and wolf genomes as above from a reconstruction of the TMRCAs from these plus one coyote. To compare the estimates from all three reconstructions, we took rolling averages across windows of 200,000 base pairs with offset of 1 bp for both the stepwise constant functions of tree heights of the reconstructed genealogies and of 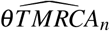.

While all three methods show different scales, we see that their general profile matches well (Figure 7), highlighting a potential to use 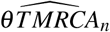 as a windowed statistic across the genome. Again, we see a tendency for 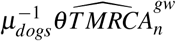 to estimate consistently lower than `tsinfer/tsdate` - a pattern that at least in our simulated data was caused by a tendency of `tsinfer/tsdate` to estimate too high TMRCAs. However, here we consider genealogies of different species, which does not mirror our single-species simulation approach necessarily.

**Figure 7:**
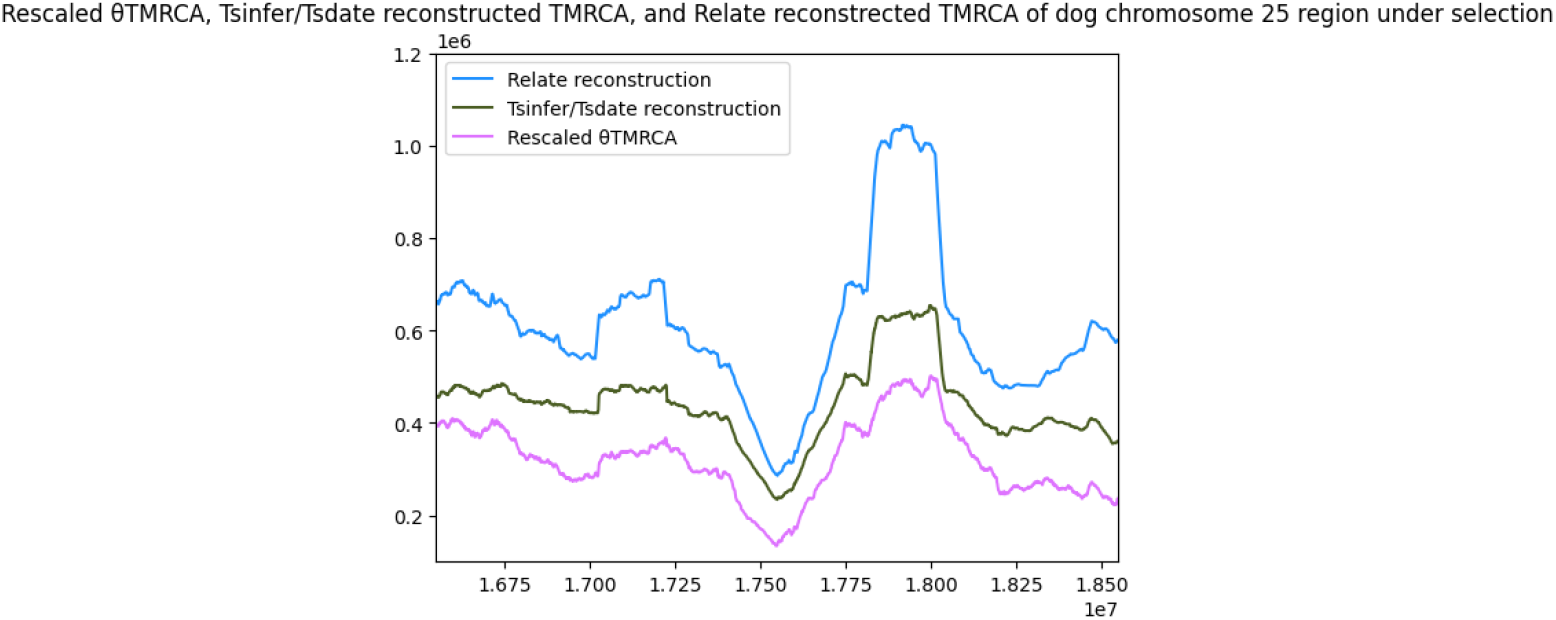
Rolling averages of tree heights in genomic windows of length 200,000 bp’s from reconstructed genealogies via `tsinfer`/`tsdate` (in black), via `Relate` (in blue), and estimating directly via 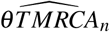, divided by *µ*_*dogs*_ · 1 (in pink).

## 7 Discussion

The simple model-free relationship (1) between the time back to the most recent common ancestor and the branch lengths spectrum leads to a surprisingly well performing estimator of the average height of ancestral trees across the genome, based on the allele frequency distribution in the sample, in other words the site frequency spectrum. This estimator, respectively a finite-sample correction by the factor *n/*(*n* − 1) has been shown to estimate the scaled mutation rate *θ* from genomic data [63] and used in the context of scanning genomic regions for selection. However, we use it here in a different context, to estimate the time to the most recent common ancestor of ancestral trees across the genome, an important property which has uses in a wide range of population genomic methods, from demography reconstruction to detecting genomic regions under selection.

The estimator returns an unbiased, but scaled moment estimate of the time to the most recent common ancestor, under a few assumptions. One of these is that the genealogical tree follows a strict molecular clock, i.e. that the mutation rate is constant across all branches of the genealogy. This assumption should typically be true when describing the ancestry of a population within a species, but may not hold if the ancestral trees are phylogenies between species. Moreover, our estimator needs point mutation counts, respective a summary of these (site frequency spectrum), thus we need to be able to call the mutant allele (polarize the site frequency spectrum).

While we consider and discuss mostly genealogies where all samples have been collected at the same time, our estimator can be extended to serially sampled population - it then measures the time from the most recent sampling time to the most recent common ancestor corrected by the average distance between sampling times and the present.

If we consider a genomic region with a single genealogy, our estimator has a straightforward interpretation as a moment estimator of the TMRCA times the total rate of mutation across this region. However, in recombining populations, genomic regions will be described by several genealogies, which typically are ancestral to an *a priori* unknown and/or inherently random fraction of this genomic region. Our estimator then is a sum of the estimators for each subregion that is described by a single genealogy. This simplifies strongly if we can assume that the mutation rate *θ*_*b*_ per basepair is constant across the genomic region and that all genealogies follow the same distribution, i.e. that if the genealogy changes, it still can be mathematically described as being generated by the same random mechanism. Biologically, this means that the ancestral lineages for different genomic regions come from comparable (comparably sized) ancestral pools and are not affected by fundamentally different evolutionary constraints (e.g., selection, introgression). In this case, 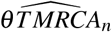 is an unbiased estimator of the expected TMRCA of the underlying genealogy model (random process) times the scaled mutation rate *θ* = *lθ*_*b*_ of the complete genomic region. For practical considerations, this should still be approximately true if a vast majority of genomic regions evolve under the same constraints, or when we are able to constrain our analysis on regions sharing the same evolutionary constraints. An example for this would be regions that evolve neutrally, in contrast to regions under selection or which are introgressed.

How precise is 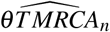 as an estimator for the expected TMRCA? To make further statistical statements, we need to specify a mathematical model of how genealogies change due to recombination. We use sequential Markovian coalescents, which approximate the full ancestral recombination graph. We do not only discuss the standard sequential Markovian coalescent whose underlying single locus genealogy is Kingman’s coalescent, but also cases where the single locus genealogies are given by multiple-merger coalescents. Multiple-merger coalescents model genome-wide diversity for a wide range of evolutionary scenarios for recombining populations - from standard Wright-Fisher type evolution without selection pressure and offspring numbers not varying strongly between individuals (described well by a Kingman coalescent genealogy) to scenarios of rapid or pervasive selection, sweepstake reproduction or recurrent extreme bottlenecks, and further scenarios. For such models, we show that the estimators are consistent for the average height of the underlying one-locus model.

Mathematical consistency is reached for genome length going to infinity. So for practical considerations, how much genome length is necessary for the estimator to be close to the expected value? While we do not mathematically assess this question here, the necessary genome size should be negatively correlated with the recombination rate, as we need have enough changes in the genealogies to reach close to the expected value (to reach a state close to ergodicity resp. well mixing - a recent result [57] considers this direction). However, we could show via simulation that, under realistic settings for (partial) human chromosomes, our method estimates the TMRCA with smaller errors than extracting TMRCAs from a state-of-the-art reconstruction method for ancestral recombination graphs, i.e. local genealogies across the considered genomic region of approximately the size of a small human chromosome.

The consistency of 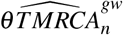, so the estimator divided by genome length, under sequential Markovian coalescents with or without multiple mergers, is a consequence of the (conditional) ergodicity of these processes. Apart from the standard sequential Markovian coalescent, we consider two types of Markovian processes with local multiple merger genealogies. The first model class considers that coalescence times of local trees are not necessarily shared across all genomic regions, but typically only in genomic regions close to each other. This model was introduced in [33] as the sequential Markovian Beta Coalescent, which we have formally extended to the full class of Ξ-coalescents. This class of geneaology models feature properties that mimick genomes under many independent selection events (similar to [16]). We can show that this Markov model, apart from the purely star-shaped coalescent and similar models, is ergodic - regardless of the starting genealogy, for long genomes the genealogical trees will start resembling a random tree coming from the underlying one-locus coalescent model.

The second class of sequential Markovian multiple merger coalescents describes models where the same processes affect all genomic regions - which we would expect for most phenomena like drastic bottlenecks, rapid selection or sweepstake reproduction ([15], [5], [2]). This is a sequential Markovian coalescent version of the models implemented in `msprime`, following the foundational concept laid out in [34]. In this second class, the genealogical trees are ergodic conditionally on the process regulating the appearance of multiple mergers. This process corresponds to the pedigree [15, 2, 42]. Biologically, this process describes the events that lead to ancestral lineages with very many offspring lines, so that ancestry of genomic material at all genomic regions may trace back to these.

This (conditional) ergodicity means that not only the TMRCA or 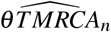 averages across a long genome towards its expectation, but also many other branch lengths- and allele frequency-based statistics. Most notably, this holds for the site frequency spectrum (SFS) - so our results provide a mathematically rigorous justification for treating the SFS in a genomic region as a sample from the genome-wide SFS, an assumption behind the sweepfinder algorithm [44, 12, 46] and used as a standing assumption elsewhere, e.g. in [11] or [23].

While our mathematical and simulation-based evidence highlight that 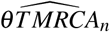 estimates the expected TMRCA well from large enough genomic regions, a reanalysis of Chromosome 25 in dogs and wolves shows a similar profile from our method and TMRCA profiles extracted from one ARG reconstruction method (`tsinfer/tsdate`), and at least a similar qualitative pattern (regions with low and high TMRCA estimates) across the region also for another method (`Relate`). As the current practice often will extract TMRCAs directly from ARG reconstructions, we have benchmarked against these methods to highlight the potential of 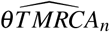 for being a useful and computationally cheap approach to estimate ancestral tree heights across the genome, especially when mostly interested in the qualitative patterns. We highlight that further TMRCA reconstruction methods exist, e.g. [32] and [65].

As our estimator is computationally cheap, it may be worthwhile to assess whether it has potential to be used within existing ARG reconstruction methods as a pivot on tree heights, or to provide a guide value for tree heights across the genome. We leave this for future work.

## 8 Data and code availability

All simulation and and inference scripts are available at https://github.com/fabfreund/thetaTMRCA. SNPs from Chromosome 25 of dogs and wolves was extracted as a subset of publicly available data accumulated in [49] - who made the full data set available as a vcf file via bioproject PRJNA448733 at NCBI: https://www.ncbi.nlm.nih.gov/bioproject/PRJNA448733.

## 9 Acknowledgements

We thank Leo Speidel for sharing the Relate analysis output for dogs and wolves with us.

ASJ acknowledges Verónica Miró Pina for preliminary discussions on the topic and DGAPA-PAPIIT-UNAM grant IN-105726 for partial support.

## 10 Appendix

### 10.1 Proof of (1) and (3)

For a formal proof of (1), we view a tree with *n* leaves as a process (*τ*_*t*_)_*t*≥0_ on the partitions of {1,…, *n*}, where we start at time *t* = 0 with blocks {1},…, {*n*} (the leaves), and the state *τ*_*t*_ consists of the partition blocks given by the leaves connected to each tree branch present at time *t* in the tree. If we then denote by *N*_*i*_(*t*) the number of blocks of size *i* at time *t* ≥ 0 and observe that

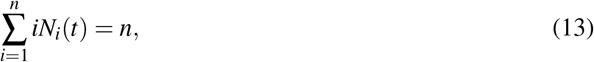

Formula (1) is obtained by integrating the latter relation between 0 and *TMRCA*_*n*_, the time the tree reaches its root.

If we consider the ancestral recombination graph, let *N*_*i j*_(*t*) be the sum of the number of blocks of size *i* at locus *a* with the number of blocks of size *j* at locus *b*. Then

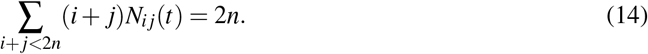

By integrating this formula between 0 and the time to the most recent common ancestor at both loci 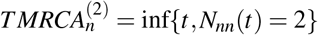, we obtain (3).

### 10.2 The serial sampling case

In the case of serial sampling, the main statistics then becomes

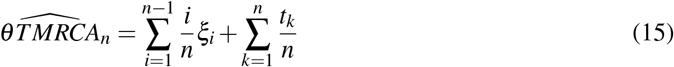

when the serial sampling times are known.

Again, under any random tree model with serial sampling at these times, the estimator is unbiased for *θ* E[*TMRCA*_*n*_] for the serially sampled *TMRCA*_*n*_. We can also define all coalescent processes for serial sampling, we can simply add new lines starting at times *t*_*k*_ to the currently present lineages. From this, both the ergodicity of the serially sampled versions of all sequential Markovian coalescents discussed as well as the consistency of the genome-wide version 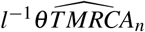 of the estimator from Eq. 15 follows analogously to the ultrametric case.

The proof is easily generalized to other types of genealogical structures as long as relations of the form (13) can be established.

#### 10.3 Proof of Proposition 3.1

Plugging (6) into (1), we get

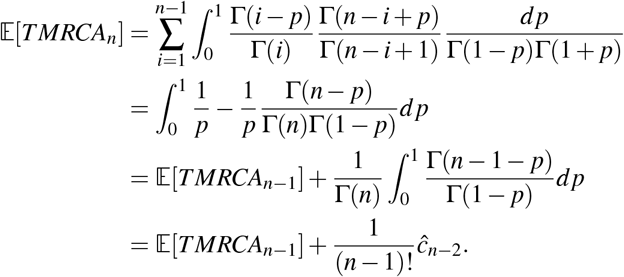

The second equality comes from Polya urns classical computations (starting configuration with one red ball of mass *p* and one blue ball of mass 1 − *p, n* − 1 draws, probability of drawing at least one red ball, the index *i* in the sum denotes the number of blue balls at the end of the experience).

#### 10.4 Poisson construction of Ξ-coalescents

Here we recall the construction of coalescents from [54]. Split Ξ = *cδ*_0_ + Ξ^′^, where *cδ*_0_ is a point mass on the zero element of Δ with weight *c* that governs purely binary collisions and Ξ^′^((0, 0, …)) = 0. To construct the Ξ-coalescent with *n* leaves, start with *n* lineages at time 0. Any present pair of lineages can merge at rate *c*, and any set of lineages present can also merge at times *t* given by atoms *p* = (*t, x*) of a Poisson point process Ψ on [0, ∞) × Δ with intensity measure 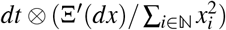, following a paintbox construction: *k*-th present lineage at time *t* generates a new, independent from all other random variables in the construction, random variable *X*_*k*_(*p*) ∼ *X* (*p*) on ℕ_0_ with *P*(*X* (*p*) = 0) = 1 − ∑_*i*∈ℕ_ *x*_*i*_ and *P*(*X* (*p*) = *i*) = *x*_*i*_. Then, all lineages *k* whose *X*_*k*_(*p*) have generated the same *i* ∈ N get merged (no mergers for lineages generating 0).

### 10.5 Proof of Theorem 5.1

We write the main steps and heuristics of the proof, avoiding tedeous notations and technicalities.

#### Invariant distribution

Consider any Ξ-*n*-coalescent. Start with a tree drawn from that distribution. We want to show that this distribution is invariant for the SMΞC. For this, it suffices to show that the transition at each of the Poisson points of the recombination-controlling Poisson process does not change the one-locus Ξ-coalescent distribution. Clearly, on the event that the Poisson point cuts no branch free, nothing changes. Now, consider the case we free a lineage via a recombination event and reglue it as described in Section 5.1 does not change this distribution. So, consider an atom (*x, t, k*), which at genomic position *x* hits the tree at time *t* on an ancestral lineage labeled *k*, involving the creation of a new tree. Suppose that the tree has *b* lineages at time *t*, from which one is freed. By the Markov property, the remaining process after time *t* is a Ξ-(*b* − 1)-coalescent, independent of the process before time *t*. The transitions described in the regluing procedure yield that the process after *t* is a Ξ-*b*-coalescent, see details in [14] that can be easily adapted to the case of Ξ-coalescents. So also on the event of cutting and regluing, the distribution does not change. Taken both cases together shows that the tree directly after genome position *x* is thus again a Ξ-*n*-coalescent.

#### Uniqueness

Suppose that Ξ has not only mass on Δ_*coll*_ or *n* = 2. We start with any tree with *n* leaves. Consider any of its mergers of size 2 ≤ *k* ≤ *n* − 1. We want to show that, after enough genomic length and recombination events, this merger is no longer present in the SMΞC almost surely. The freeing and regluing procedure implies that this merger can decrease by one or increase by one with strictly positive probabilities. If the merger size is *n*, it decreases by one with strictly positive probability. If the merger size is 2 and it decreases by one, it then disappears. We then start building up new purely binary and binary multiple merger coalescent event with the rates given in the construction, the latter than can be extended into multiple mergers - and this construction does not depend on the starting state.

#### Birkhoff Theorem

Equation (1.2) in [51] shows (10). From the two previous steps, and following Theorem 20.10 in [27], we obtain that the SMΞC is strongly ergodic, and any starting tree distribution at genomic position 0 converges to the invariant distribution. The TMRCA of the coalescent tree does have finite *p*th moments for *p* ≥ 1 under the invariant distribution. This follows from seeing that the waiting time for the next coalescence in any state of the Ξ-*n*-coalescent is stochastically smaller than an exponential random variable with a finite parameter depending on Ξ. Finally, the relation (1) and the *p*-integrability of the TMRCA imply the *p*-integrability of the length vector (*L*_1_, …, *L*_*n*−1_).

##### Remark 10.1.

Ξ*-coalescents with n* ≥ 3 *having all the mass of* Ξ *in* Δ_*n*_ *are not ergodic. Note that these are coalescents with no purely binary mergers. To see that they are not ergodic, consider a genealogy at genomic position 0. Consider a tree whose first coalescence event c*_1_ *is non-binary at time t*_1_ *(this is guaranteed if we start with a tree that is a realisation of a* Ξ*-coalescent with such a* Ξ*). Then, the structure of the rates for collisions given by [54, Eq. (11)] for such measures* Ξ *guarantees that i) there cannot be any binary mergers before c*_1_ *[the rate of a binary merger is 0, as it corresponds throwing n balls in n* − 2 *boxes, but only 2 would land in the same box, which is not possible] and ii) the merger at t*_1_ *cannot be fully dissolved, but only reduced to at least triple (or bigger) merger respectively a simultaneous double binary merger. To see the latter, recall that the rates appearing in the regluing operation of a freed lineage correspond to each of n lineages throw a ball at at most n* − 2 *boxes, conditional on all merged lineages already have landed in the same box(es, for simultaneous mergers multiple boxes). While this can lead to lineages not rejoining, after several of this free-and-not-rejoining events in case let’s assume we would break up a last triple merger or double binary merger. This would correspond, in the balls-in-boxes interpretation of the transition rates, to that only 2 balls would land in the same box and thus merge, but that the remaining n-2 balls would not not merge at this time, so would need to land in separate boxes. However, there are only n* − 3 *separate boxes at most apart from the box leading to the already present merger. Hence, this is not possible. So we will again have at least a triple or double binary merger at t*_1_, *and cannot dissolve this merger any further - in each tree across the genome there will be a merger at time t*_1_. *Hence, we cannot asymptote towards the one-locus* Ξ*-coalescent distribution across the genome, which is a property of ergodicity*.

*An example for such coalescent are star-shaped coalescents (*Ξ *is a point measure in* (1, 0, 0, …)*) where all present lineages merge after an exponential waiting time*.

### 10.6 Proof of Corollary 5.1

For each *i*, the genome-wide entry *ξ*_*i*_ is the sum of independent Poisson random variables on the branches subtending *i* leaves for each tree in the SMΞC. By denoting such length at genomic position *l*^′^ by *L*_*i*_(*l*^′^), we have that the conditional law of *ξ* is Poisson with parameter 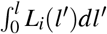. If we then consider the Laplace transform of *l*^−1^*ξ*_*i*_, we see, for *λ >* 0,

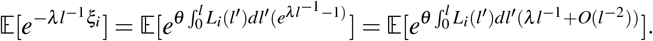

By Theorem 5.1, this quantity converges to 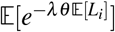 when *l* → ∞. We used the series expansion for the inner exponential function. This establishes convergence to the constant *θ* E[*L*_*i*_] in distribution, and hence in probability. This holds for all *i*, and then (1) establishes the proof.

### 10.7 Proof of Theorem 5.2

The general proof strategy is analogous to proving Theorem 5.1, but conditionally on Ψ = *ψ*. Conditioning on Ψ only means that instead of random Poisson points, we have deterministic points *ψ*. We first show that the distribution of the tree at position 0 in the genome (which follows the Poisson construction, but with a fixed realization of the Poisson point process) is not changed by the freeing-and-regluing operation. We will condition on Ψ = *ψ* for the rest of the proof.

By freeing a lineage *k* in the Ψ-directed SMΞC at time *t*_*r*_, to determine its mergers with other present lineages, we simply switch its variables *X*_*k*_(*p*), for all *p* = (*t, k*) ∈ *ψ* with *t > t*_*r*_, to copies that are independent of all other variables - which leads to an identical probabilistic structure as before the freeing, as the behavior at the Poisson points was also controlled by identical and independent *X*_*k*_(*p*) with the same distributions. Additionally, any binary collisions appearing with rate *c* are again distributed exactly as before the freeing of lineage *k* by definition. Thus, an invariant distribution is given by the Ξ-coalescent conditioned on the realization of the Poisson process (the quenched coalescent from this setup).

Next, we show that this invariant distribution is unique by arguing (again as in Theorem 5.1) that any two starting states (different tree and paintbox variables, but identical realisation Ψ = *ψ*) will run into the same distribution for genomic position *l* → ∞. As the mergers of any freed lineage *k* are completely governed by their paintbox variables (*X*_*k*_(*p*))_*p*∈*ψ*_, we only need to observe that any lineage will almost surely be freed at an arbitrarily small time -thus replacing the original two starting states with independent copies from the same distribution of all merger variables. as well as new binary mergers occurring with rate *c* between any present pairs. The distribution at this point is thus independent from the two starting states, and is, by construction, the invariant distribution we found earlier.

As the conditional Ψ-directed SMΞC is Markovian, we can again use the Birkhoff theorem [51] to prove (10). From our results of *p*-integrability of the TMRCA, it follows for almost all realizations of the Poisson point process that the TMRCA is *p*-integrable also in the conditioned version of the one-locus coalescent, and the *p*-integrability of other branch lengths follows as in the proof of Theorem 5.1. Finally, Corollary 5.1 also holds conditionally for the Ψ-directed SMΞC with the same arguments.

#### Remark 10.2.

*By construction, we assure that conditioning on the state of the Poisson process* Ψ *controlling the multiple mergers across the genome leads to a regular conditional distribution, and this is used implicitly in the proof, where we show ergodicity conditional on* Ψ. *Moreover, while we are mainly, for biological application, interested in the result conditioned on a specific realisation of* Ψ, *we can also formulate, by integration over the regular conditional distribution conditional on* Ψ, *the corresponding unconditional result. For any L*^*p*^*-integrable function f of the* Ψ*-controlled sequential coalescent* (*C*_*g*_)_*g*∈[0,∞)_ *we almost surely have*

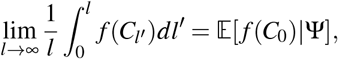

*where the latter is a quenched coalescent*.

## Notes

### Competing Interest Statement

The authors have declared no competing interest.

